# Index hopping on the Illumina HiseqX platform and its consequences for ancient DNA studies

**DOI:** 10.1101/179028

**Authors:** Tom van der Valk, Francesco Vezzi, Mattias Ormestad, Love Dalén, Katerina Guschanski

## Abstract

The high-throughput capacities of the Illumina sequencing platforms and the possibility to label samples individually have encouraged a wide use of sample multiplexing. However, this practice results in read misassignment (usually <1%) across samples sequenced on the same lane. Alarmingly high rates of read misassignment of up to 10% were reported for the latest generation of lllumina sequencing machines. This may make future use of the newest generation of platforms prohibitive, particularly in studies that rely on low quantity and quality samples, such as historical and archaeological specimens. Here, we rely on barcodes, short sequences that are ligated to both ends of the DNA insert, to directly quantify the rate of index hopping in 100-year old museum-preserved gorilla (*Gorilla beringei*) samples. Correcting for multiple sources of noise, we identify on average 0.470% of reads containing a hopped index. We show that sample-specific quantity of misassigned reads depend on the number of reads that any given sample contributes to the total sequencing pool, so that samples with few sequenced reads receive the greatest proportion of misassigned reads. Ancient DNA samples are particularly affected, since they often differ widely in endogenous content. Through extensive simulations we show that even low index-hopping rates lead to biases in ancient DNA studies when multiplexing samples with different quantities of input material.

## 1| Introduction

Multiplexing samples for sequencing is common practice in genomic studies (Craig *et al*. 2008; Meyer & Kircher 2010; Smith *et al*. 2010). During multiplexing, samples are individually labelled with unique identifiers (indices) that are embedded within one (single indexing) or both (dual indexing) sequencing platform-specific adapters (Meyer & Kircher 2010; Kircher *et al*. 2012). The samples are subsequently pooled into a single DNA library and sequenced on the same lane, greatly reducing per sample sequencing cost. Following sequencing, computational demultiplexing based on the sample-specific indices enables the assignment of sequenced reads to the respective sample of origin. In recent years, the output from sequencing platforms has increased dramatically, making multiplexing the recommended standard sequencing workflow on the latest generation of Illumina platforms (e. g. NovaSeq) (Illumina Inc.). However, ever since multiplexing approaches were introduced, low rates of read misassignment across samples sequenced on the same lane have been reported on all Illumina platforms (Kircher *et al*. 2012; Nelson *et al*. 2014; Renaud *et al*. 2015; Wright & Vetsigian 2016; D’Amore *et al*. 2016). Read misassignment is the result of reads carrying an unintended index and consequently being erroneously attributed to the wrong sample. Processes resulting in read misassignment, i.e. presence of reads with an incorrect index, are numerous. The effect of sequencing errors that can convert one index sequence into another is well known and has led to series of recommendations for designing highly distinct indices (Meyer & Kircher 2010). Jumping PCR during bulk amplification of library molecules that carry different indices can generate chimeric sequences and should be avoided (Meyerhans *et al*. 1990; Odelberg *et al*. 1995; Carlsen *et al*. 2012; Esling *et al*. 2015). Similarly, cross-contamination of indexing adapters during oligonucleotide synthesis or laboratory work can lead to reads obtaining an unintended index. Additionally, cluster misidentification due to “bleeding” of indices into neighbouring clusters have been reported on all high throughput sequencing platforms (Kircher *et al*. 2012; Nelson *et al*. 2014; Renaud *et al*. 2015; Mitra *et al*. 2015; Wright & Vetsigian 2016; D’Amore *et al*. 2016; Vodák *et al*. 2018). For the latest Illumina platforms with patterned flow cells and ExAmp chemistry (e.g. Hiseq X and NovaSeq), it has been suggested that read misassignment is caused by the presence of free-floating indexing primers in the final sequencing library (Illumina Inc. 2017; Sinha *et al*. 2017). Such free-floating molecules can appear if sequencing libraries are not stored properly and become fragmented, or if the final sequencing libraries retain non-ligated indexing primers due to inefficient clean-up and size selection (Illumina Inc. 2017). These free-floating primers can anneal to the pooled library molecules and become extended by the DNA polymerase before the rapid exclusion amplification on the flow cell, creating a new library molecule with an erroneous index (Figure 1). We refer to this particular process of generating misassigned reads as **index hopping.** The reported rate of read misassignment on Illumina platforms that rely on the traditional bridge amplification for cluster generation is low (<1%) (Kircher *et al*. 2012; Nelson *et al*. 2014; Wright & Vetsigian 2016) and therefore this source of error has been readily ignored. However, on the latest Illumina patterned flow cell platforms with ExAmp chemistry, the reported rate of read misassignment ranges from 0% to 10% (Illumina Inc. 2017; Sinha *et al*. 2017; Owens *et al*. 2018; Griffiths *et al*. 2018; Vodák *et al*. 2018), with Illumina quoting a read misassignment rate of up to 2% (Illumina Inc. 2017).

**Figure 1:**
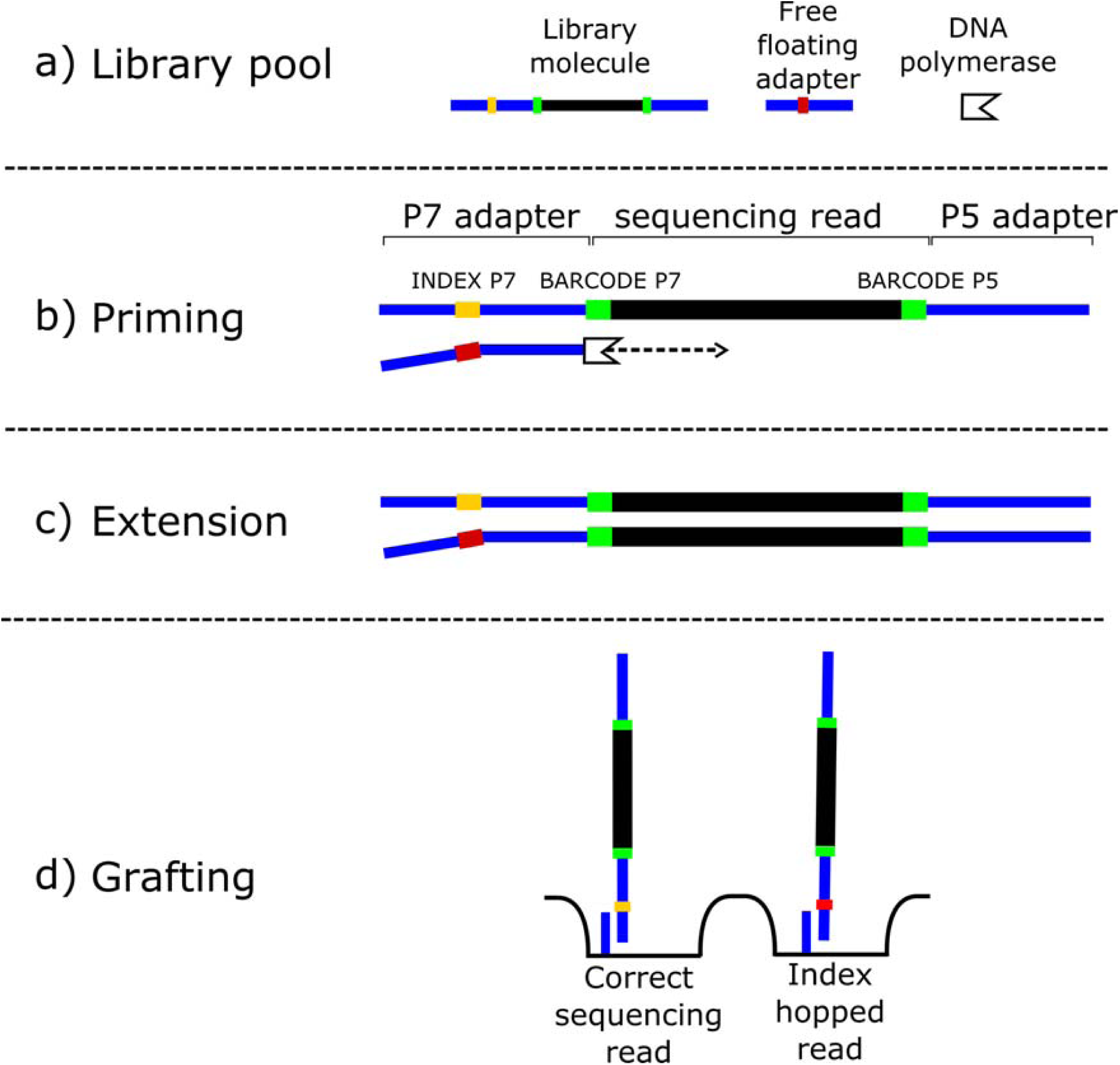
Schematic of index hopping during ExAmp clustering. A) The library pool, containing barcoded and indexed library molecules (black: DNA insert, green: P5 and P7 barcodes, orange: P7 index) and free-floating indexing primers, is mixed with ExAmp reagents before loading on the patterned flow cell. B) Free-floating adapter anneals to the adapter sequence of a library molecule and C) the library molecule gets extended by the DNA polymerase, forming a new library molecule with a wrong index. D) The library molecules are denatured, separating the strands, and each library molecule is allowed to graft onto a nanowell on the patterned flow cell.

As a consequence of conflicting results, the prevalence and severity of read misassignment on the latest Illumina platforms remain unclear. This is partly due to the difficulties of reliably identifying misassigned reads in sequencing experiments, particularly when pooling similar samples types (e.g. multiple individuals from the same population that have high sequence similarity). The use of dual indexing allows for the filtering of the majority of reads that show signs of read misassignment (Kircher *et al*. 2012). However, since indices can potentially be switched at both ends of the molecule and the number of available indices is limited, it remains difficult to directly quantify read misassignment rates on these platforms. Consequently, the so far reported rates off index-hopping have been estimated using indirect methods (Sinha *et al*. 2017; Owens *et al*. 2018; Griffiths *et al*. 2018; Vodák *et al*. 2018).

The reported high rates of index-hopping are especially worrisome for studies involving sequencing data obtained from degraded samples such as ancient and historical specimens, since in most cases such studies rely on low-coverage genomic data (Shapiro & Hofreiter 2014). Inferences are therefore often based on subtle differences between limited sets of polymorphic sites, so that even small quantities of misassigned sequencing reads can potentially lead to erroneous conclusions. It is thus crucial to distinguish genuine sample-derived endogenous DNA fragments from false signals (Skoglund *et al*. 2014).

The purpose of this study is two-fold. First, we aim to directly quantify the rate of index-hopping on the Illumina patterned flow cell platforms for a standard ancient DNA library. To this end, we make use of inline barcodes, short unique seven base pair sequences ligated to both ends of the DNA fragments (Rohland & Reich 2012), in combination with the indexed primers that are traditionally used for sample identification. The barcodes become part of the sequencing read and thus allow for accurate identification of the read origin, even in the presence of index hopping. Therefore, the amount of index hopping can be directly quantified by identifying reads with wrong barcode-index combinations. Second, we aim to identify and characterize biases resulting from index-hopping in pooled ancient DNA libraries that may impact downstream analyses typical for ancient DNA research. To achieve this, we simulate ancient DNA sequencing libraries under different rates of index-hopping and quantify the impact of misassigned reads on population genomic inferences by performing a set of standard genome-wide analyses.

## 2| Methods

### 2.1 | Library preparation

DNA extracts from seven historical eastern gorilla (*Gorilla beringei*) samples were turned into sequencing libraries as described in (van der Valk *et al*. 2018) (see supplementary material). All library preparation steps except indexing PCR were performed in a dedicated ancient DNA laboratory to minimize contamination. Briefly, 20 μl DNA extract was used in a 50 μl blunting reaction together with USER enzyme treatment to remove uracil bases resulting from aDNA damage (Briggs *et al*. 2010). DNA fragments within each sample were then ligated to a unique combination of incomplete, partially double-stranded P5- and P7-adapters, each containing a unique seven base pair sequence (Rohland *et al*. 2015) (Table S1). We refer to these as the P5 and P7 **barcodes** from here on. All barcode sequences were at least three nucleotides apart from each other to ensure high certainty during demultiplexing and avoid converting one barcode into another through sequencing errors (Rohland *et al*. 2015) (Table S1). To increase the complexity of the pooled sequencing library, one sample (sample 7) was split in two fractions, each of which received a different barcode combination (Table 1).

**Table 1:** Sequencing statistics and estimates of contamination and index hopping.

| *Sequencing run* | *Sequencing reads after quality filtering* | *Sequencing reads with wrong barcode pairs* | *Sequencing reads with wrong barcode pairs (%)* | *Sequencing reads with wrong index-barcode combination* | *Sequencing reads with wrong index-barcode combination (%)* |
| --- | --- | --- | --- | --- | --- |
| *Run 1* | 316203540 | 99575 | 0.0301 | 1543241 | 0.488 |
| *Run 2* | 127766205 | 42457 | 0.0315 | 837926 | 0.656 |
| *Run 3* | 429511898 | 94527 | 0.0211 | 2740612 | 0.638 |
| *Average* | 291160548 | 78853 | 0.028 | 1707260 | **0.594** |

Indexing PCR was performed for 10 cycles using a unique P7 indexing primer for each sample, as in (Meyer & Kircher 2010) (Table S1). We refer to the unique sequence added during the indexing PCR as the P7 **index.** As with the barcodes, all index sequences differed by at least three base pairs from each other (Table S1). Indexing PCR for sample 7 was performed in a single reaction combining both fractions of this sample. Following the indexing PCR, each DNA fragment contained three unique identifiers: the P5 and P7 barcodes directly ligated to the ends of the DNA fragments, and the P7 index contained within the Illumina sequencing adapter (Figure 1). Sample libraries were cleaned using MinElute spin columns, fragment length distribution and concentrations were measured on the Bioanalyzer. We then pooled all seven sample libraries in a ratio of 2:1:2:1:1:1:2 for samples 1 to 7, and performed two rounds of AMPure XP bead clean-up, using 0.5X and 1.8X bead:DNA ratio,respectively. We confirmed that indexing primers were successfully removed during clean-up by running the final library on a Bioanalyzer (Figure S1). The pooled library was sequenced on three HiSeqX lanes that were part of independent runs with a 5% phiX spike-in, at the Science for Life Laboratory in Stockholm.

### 2.2 | Data processing

All reads were demultiplexed based on their unique indices using Illumina’s bcl2fastq (v2.17.1) software with defaults settings, allowing for one mismatch per index and only retaining “pass filter” reads (Illumina Inc.). All unidentified reads, i.e. reads containing indices not used in our experiment, were retained and subjected to the same filtering steps as assigned reads (see below). We removed adapter sequences using AdapterRemoval V2.1.7 with standard parameters (Schubert *et al*. 2016). Due to the fragmented nature of DNA in historical samples, we could subsequently merge the reads, requiring a minimal overlap of 11bp and allowing for a 10% error rate. The merging of reads allowed us to obtain sequencing information for the complete DNA molecule and thus to accurately identify the barcodes on both ends of the DNA fragment (P5 and P7 barcodes, respectively, Figure 1). Unmerged reads and reads shorter than 29 basepairs were removed. To increase certainty, we only retained reads with error-free P5 and P7 barcodes and an average quality score of at least 30 using prinseq V0.20.4 (Schmieder & Edwards 2011).

### 2.3 | Disentangling cross-contamination from index hopping

Low rates of cross-contamination of barcodes and indexes can be expected, even if strict measure are followed during library preparation, such as the use of clean-room facilities (Kircher *et al*. 2012). This can result in reads containing a wrong index-barcode pair and could be falsely interpreted as evidence for index hopping. Since the inline barcodes used in this experiment are unaffected by index-hopping (Figure 1), we can accurately estimate the rate of barcode cross-contamination as the fraction of reads containing a P5-P7 barcode pair that was not used during library preparation. In rare cases, barcode cross-contamination results in a read with a valid barcode pair (e.g. a barcode combination that was intendedly used during library preparation) and thereby remain undetected in our estimate. However, since we used every barcode only once, the proportion of reads resulting from such an event is several orders of magnitude lower than the fraction of reads containing an invalid barcode pair and does therefore not significantly affect any of our estimates (see supplementary material).

As the Illumina HiSeq X platform did not support a dual-indexing design at the time of this experiment, the rate of index cross-contamination could not be estimated using invalid index pairs. Therefore, we relied on the fact that of the 40 indices that are routinely used in our laboratory only seven were implemented in this experiment (Table S2). Assuming a relatively equal rate of cross-contamination between all 40 indexes, we estimated index cross-contamination as the fraction of reads containing any of the 33 indices that were not deliberately included during our experiment.

We then determined the raw rate of index hopping as the fraction of reads showing an index-barcode combination not used during the library preparation. We accounted for the possibility of barcode and index cross-contamination resulting in the same barcode-index combination by subtracting the contamination estimates obtained above from the raw value of index hopping. All statistical analyses were performed in R 2.15.3 (Team R Core 2016) (see supplementary material).

### 2.4 | Simulations of aDNA sequence libraries

To quantify downstream biases resulting from index-hopping during pooled sequencing, we simulated ancient DNA sequencing libraries with different endogenous content under varying rates of index hopping. First, four “template” genomes were simulated to serve as seeds for four populations, popA, popB, popC and popD, respectively, by using chromosome 1 of the gorilla reference (removing all N nucleotides) (Gordon *et al*. 2016). The population divergence was set as follows: popA – popB: 20.000 years, popC – popD: 20.000 years and popA/popB – popC/popD: 200.000 years (Figure 3b) and we introduced random mutations at a rate of 1.67 · 10^-9^ per base per year (corresponding to the estimated gorilla mutation rate (Besenbacher *et al*. 2018)). We then simulated thirty individuals for each population, using the “template” genomes as a starting point and introducing on average 5 · 10^-7^ random mutations per base (corresponding to all individuals within each population sharing a common ancestor on average 300 years ago). We did not simulate any admixture between the populations. Next, each individual genome was converted into an ancient DNA sequence library (fastq-format), with insert size normally distributed around 50bp and endogenous content of either 0.133%, 0.398%, 1.33%, 3.98%, 13.26% or 39.78% to mimic characteristics often observed in ancient DNA studies. The levels of endogenous content were chosen to result in commonly observed genome coverages of ancient DNA samples (see below). The non-endogenous reads consisted of fastq-reads simulated using the PhiX-reference genome (NCBI nucleotide ID: NC_001422) as template. We then simulated sequencing output of equimolar pooled libraries as would be obtained from sequencing the pools on four NovaSeq6000 runs (flow cell-type S4, expected output 8-10 billion reads per run). The expected output of ~40 billion reads thus consisted of a random sample of ~333 million fastq-reads from each of the 120 simulated sequencing libraries (30 individuals x 4 populations). Index-hopping was simulated by giving each read a predefined probability of randomly hopping into another sample, using the following rates: 0.0%, 0.1%, 0.5%, 1%, 5% and 10%, reflecting the levels of index-hopping reported in the literature. We did not simulate indels/deletions, PCR duplications, sequencing errors, and post-mortem DNA damage, since we specifically aimed to address the biases resulting from index-hopping. Our final simulated data thus consisted of six datasets of 120 simulated ancient DNA libraries with varying endogenous content obtained from four different populations, with a different level of index-hopping in each of the six datasets.

To analyse the simulated data, we aligned all reads per individual to the gorilla reference chromosome 1 (Gordon *et al*. 2016) (note that in our simulations this reference represents the ancestral state for each site) using bwa-mem on default parameters (Li & Durbin 2009). The obtained coverage for the simulated individuals was 0.1X, 0.3X, 1X, 3X, 10X or 30X, depending on the sample’s endogenous content (note that the levels of endogenous content were chosen to result in these coverages). Next, we employed a pipeline specifically designed for analysing low-coverage genomes from degraded DNA sources. We obtained genotype likelihoods for each individual using angsd (Korneliussen *et al*. 2014), filtering reads below mapping quality of 30 (-minMapQ 30), a flag above 255 (-remove_bads 1) and removing reads with multiple hits (-uniqueOnly 1). We then only considered genotypes with a likelihood ratio test statistic of minimum 24 (-SNP_pval 2e-6) using the samtools genotype model (-GL 1).

### 2.5 | Inferring population genomic statistics from simulated data

We used Principal Components Analysis, Admixture and D-stats (ABBA-BABA test) to reconstruct population divergence under different levels of index-hopping. Principal Components Analysis was run using PCAngsd with default parameters and 200 EM iterations for computing the population allele frequencies (Fumagalli *et al*. 2013). Individual admixture proportion were obtained using NgsAdmix (Skotte *et al*. 2013) at default parameters and using K = 4 (number of ancestral clusters). Pairwise D-stats of the format (popA,popB,popC,ancesteral) were calculated for each possible pair of individuals in popA and popB by sampling a random allele at each site (htsbox pileup -R -q 30 -Q 30 -s 1, https://github.com/lh3/htsbox), using a high coverage (30X) individual from popC as the third ingroup and the ancestral allele (reference allele) at each site as the outgroup. Standard-deviations and resulting Z-scores (the number of standard deviations of D from 0) were obtained using a jackknife approach with blocksize of 2Mb.

## 3| Results

### 3.1 | Empirical data

#### 3.1.1 | Barcode and index cross-contamination

Since our sequencing libraries were made from degraded historical samples and thus contained a large proportion of short DNA fragments (Figure S1) the majority of reads could be confidently merged for all three sequencing runs (95.3% SE ± 1.0%). This allowed us to accurately infer both barcodes at the read ends. After all filtering steps (Methods), the final dataset contained 89.3% SE ± 1.9% of the original sequence reads.

We estimate the average level of barcode cross-contamination across all three runs at 0.0276% SE ± 0.0026 (see methods, Table 1, Table S3, Figure S2), with different rates observed between samples (global chi-square test, P<10^-15^). Assuming that adapter ligation of barcodes is unbiased with respect to the barcode sequence (Rohland *et al*. 2015), this low percentage of cross-contamination will lead to a negligible fraction of reads (1.09 · 10^-8^ %, see supplementary material) with a barcode pair that wrongly appear as having undergone index hopping. The rate of index cross-contamination was estimated at 0.124% SE ± 0.0023 (Table S4), by quantifying the fraction of reads containing indices that were not intentionally used in our experiment (see Methods, Table S2).

#### 3.1.2 | The rate of index hopping

Index hopping will not affect the barcodes that are directly ligated to the DNA fragments. Therefore, it can be readily distinguished from barcode cross-contamination by identifying reads containing an incorrect combination between an index and a barcode pair. Across all three sequencing runs, we detected a low proportion of such reads (mean=0.594%, SE ± 0.0434%, Table 1). However, we estimate that ~0.124% of these reads are a result of index and barcode cross-contamination (see above). Therefore, the corrected rate of index hopping in our experiment across all three sequencing runs is ~0.470% SE ± 0.044. The proportion of hopped reads differed significantly among samples (chi-square test, P<10^-15^) and was positively correlated with the number of sequenced reads per sample (Pearson’s r = 0.96, P = 0.0005, Figure 2). This suggests that in multiplexed sequencing runs, the samples with higher number of sequenced reads will serve as the dominant source of hopped reads. Even though the overall rate of index hopping is low, samples with proportionally few sequenced reads are thus considerably more affected by index hopping. In our experiments, this resulted in 2.49% SE ± 0.29% of hopped reads in the sample with the lowest number of sequenced reads (Table S4, S5, Figure 2).

**Figure 2:**
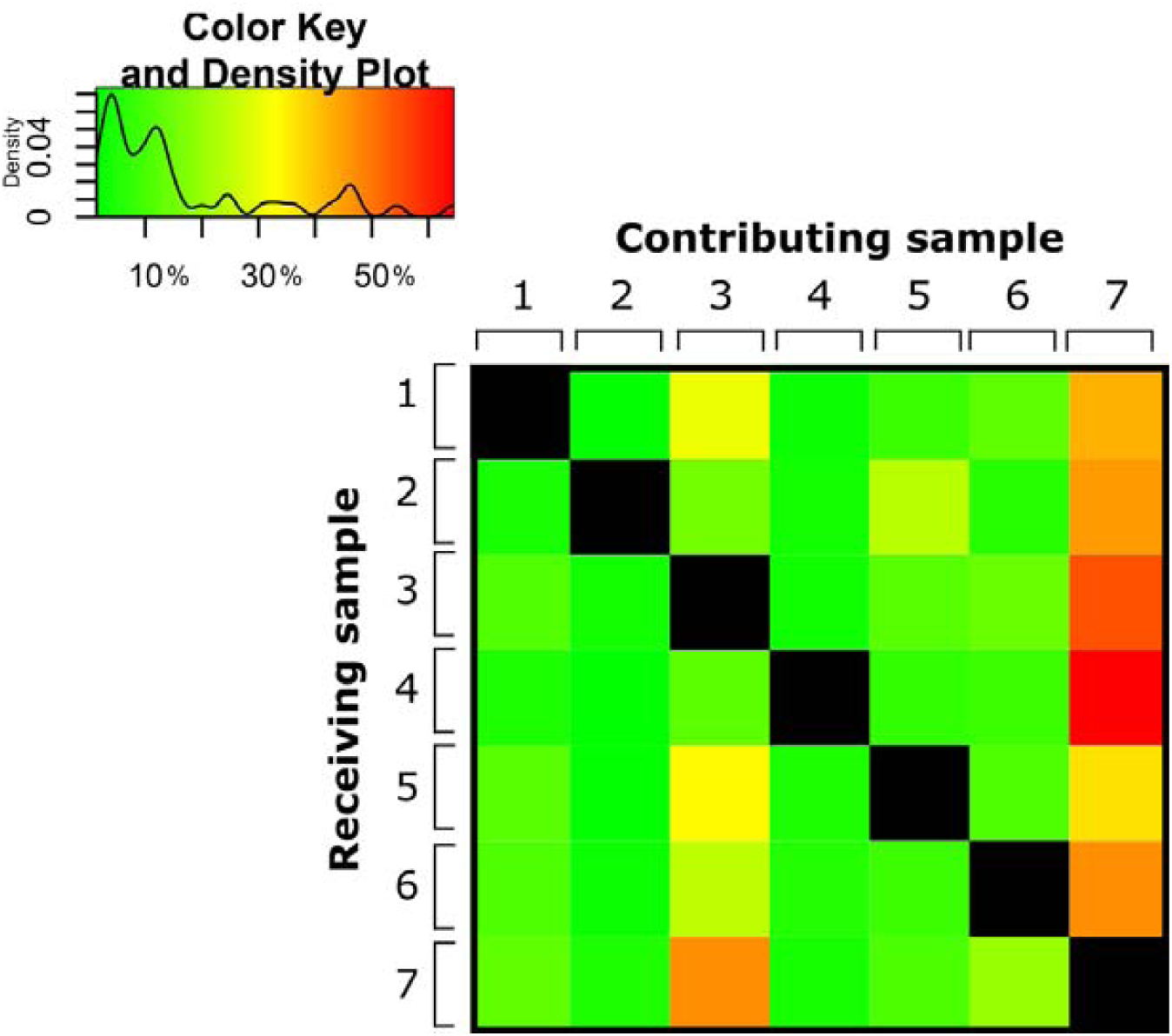
Proportion of hopped reads per sample out of all hopped reads. Samples in the top row contribute hopped reads, whereas samples on the left receive hopped reads. Samples with high number of sequenced reads (e.g. 3 and 7) are also the main contributors of hopped reads.

We find that the rate of index hopping differed significantly by read length and slightly by GC content (chi-square test, both P<10^-15^, Figure S3). Reads shorter than 90 bp and reads with GC content above 40% showed significantly higher proportion of hopped reads than expected under a random distribution.

### 3.2 | Simulated data

#### 3.2.1 | Effects of index hopping on estimates of sample endogenous content

Ancient DNA studies frequently rely on the screening of a large number of samples by means of pooled low depth sequencing to identify samples with good DNA preservation and high endogenous content. Through introduction of endogenous reads into low-quantity samples, index-hopping can lead to a false signal of DNA preservation. We estimated the endogenous content for each of our simulated ancient DNA sequencing libraries as the fraction of reads that mapped to the reference genome under different rates of index-hopping (see Methods). We observed that already at low rates of index hopping (<1%), the endogenous content of low-quality samples (0.1%-0.4% endogenous content) was over-estimated (up to ~2-fold higher, Figure 3a). This bias became more pronounced as rates of simulated index hopping increased and resulted in up to 8-fold higher estimate of endogenous content. Estimates for samples with higher endogenous content (>3%) were biased only at high rates of index-hopping (5-10%).

**Figure 3:**
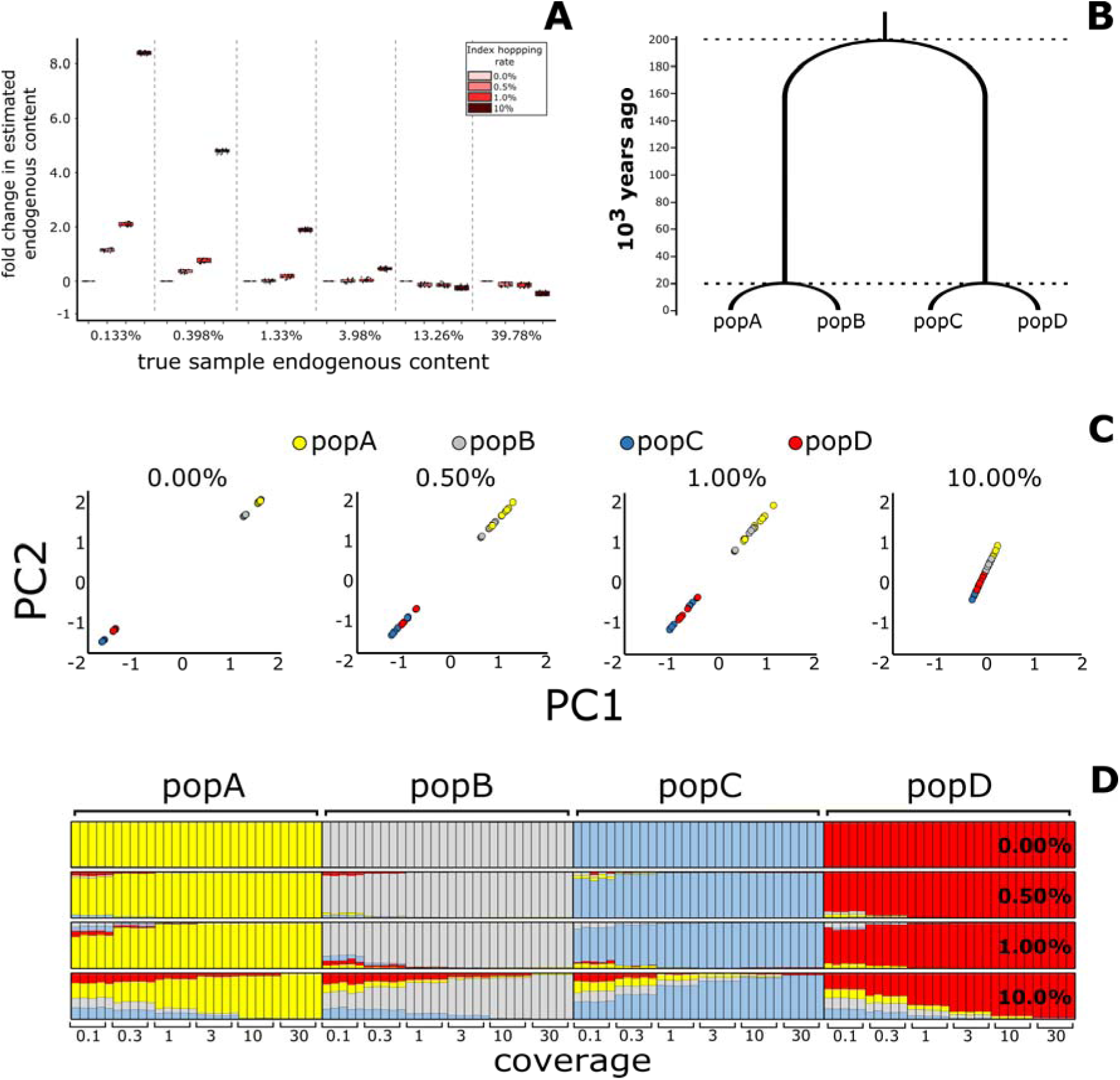
(A) Index hopping biases estimates of endogenous content. X-axis shows the simulated sample endogenous content, Y-axis shows the fold-change in the inferred endogenous content (fraction of mapped reads). Colours depict the simulated rate of index-hopping, which increases from left to right for each sample. Biases for samples with high endogenous content are minor (note that these samples appear to “loose” reads that are being assigned to samples of low endogenous content). However, samples of low quality (low endogenous content) are disproportionally affected by index hopping, leading to erroneously high estimates of endogenous DNA content. (B) Schematic representation of the simulated populations and their divergence times. For each populations 30 individuals with different levels of endogenous content are simulated. (C) Effects of index hopping on inferences of population differentiation: Principal Components Analysis for all individuals under different rates of index hopping (depicted on the top). Each plot shows all 120 individuals from the four simulated populations. (D) Effects of index hopping on inferences of individual admixture proportion. Each bar is an individual, X-axis depicts sample coverage. Percentage at the right depict simulated index-hopping rate.

#### 3.2.2 | Index hopping biases population genetic inferences

Principal Components Analysis clearly differentiates the four simulated populations from each other in the absence of index-hopping (Figure 3c, S4). However, we find that already at a low rate of index-hopping (0.50%, similar to observed in our empirical data), population differences between popA-popB and popC-popD start to disappear. This is caused by the relatively high number of hopped, wrongly assigned reads in samples with low endogenous content. At extreme levels of index hopping (10%), even the differences between the highly diverged populations (popA-popB vs popC-popD) disappears (Figure 3c, S4).

Admixture analysis corroborated the results obtained from PCA. We find that at low rates of index-hopping (0.50%), false signals of shared ancestry between individuals from different populations start to appear in the low-coverage samples (Figure 3d, S5). At the highest rate of index-hopping (10%), only the highest quality samples (e.g. 30X) remain unbiased (Figure 3d, S5).

We used ABBA-BABA counts to test if index hopping can lead to erroneous inferences of gene-flow between the populations. Although the Z-scores become skewed at low rates of index-hopping (<1%) if samples differ strongly in endogenous content, we only inferred significant deviations from zero at high rates of index-hopping (>5%) (Figure S6).

## 4| Discussion

Using a dual barcoding strategy during library preparation, we show that index hopping occurs on the Illumina HiSeq X platform, but its rate is low in our ancient DNA library (0.470% SE ± 0.044). Although multiple sources of error such as jumping PCR, barcode and index cross-contamination, sequencing errors, and index hopping can result in read misassignment, our experimental design allowed us to systematically address each of them. Jumping PCR can be eliminated as explanation for wrong index-barcode combinations, since we avoided amplification of pooled libraries from different samples. However, we show the strong effect of jumping PCR when looking at the rate of wrong barcode combinations in the only sample with two different barcode pairs that was amplified in a single indexing reaction (Fig. S2). We further show that the rate of barcode and index cross-contamination is very low (0.027% SE ± 0.0026 and 0.124% ± 0.0023, respectively) and therefore not the primary cause of observed reads with the wrong index-barcode pairs.

Read misassignment is not a novel phenomenon on the Illumina sequencing platforms. Reported error rates range from 0.1% to 0.582% on the HiSeq 2500 (Kircher *et al*. 2012; Wright & Vetsigian 2016) and from 0.06% to 0.51% on the MiSeq platforms (Nelson *et al*. 2014; Renaud *et al*. 2015; D’Amore *et al*. 2016). It is therefore noteworthy that the fraction of hopped reads as estimated in our study (0.470%) is similar to that reported for other platforms. However, it markedly differs from previous estimates for the Illumina platforms with ExAmp chemistry, which are based on sequencing modern (high quality) DNA and range from 0% to 2.5%-10% (Sinha *et al*. 2017; Owens *et al*. 2018; Griffiths *et al*. 2018). Since the sequencing chemistry of the Illumina NovaSeq platform is identical to that used for the HiSeq X, this platform is likely to be affected at a similar rate as reported here.

We used a standard library preparation protocol for degraded samples, which includes rigorous removal of free-floating adapters through size selection and cleaning (supplementary methods). This practice likely resulted in the relative low rate of index hopping in our experiment. As previously suggested, strict library clean-up and size selection is thus recommended for multiplexed ancient DNA sequencing studies.

A so far neglected observation is that the number of hopped reads into each sample is proportional to the total number of reads contributed by this sample to the pooled sequencing library. Pooling samples in different quantities leads to a greater proportion of hopped reads into samples with fewer sequenced reads. In this study, libraries with the lowest number of sequenced reads displayed up to 3.2% of misassigned reads (Table S5), an order of magnitude higher than the average rate within a lane. The effect of this skewed rate of index hopping becomes even more severe if the endogenous content is markedly different between samples. Since the endogenous content is usually not known beforehand, pooling samples in equimolar quantities can lead to large differences in the number of endogenous reads between samples. In such cases, even at the rate of index-hopping reported here, the proportion of false assigned endogenous reads within low quantity samples can reach rates above 10% (Figure S7), resulting in highly overestimated sample endogenous content (Figure 3a). This is problematic, as presence of even few reads of interest can lead to further processing and deep sequencing of an ancient DNA sample deemed to be of importance. Additionally, we detected a higher rate of index hopping among shorter reads and small differences in the fraction of index-hopped reads related to read GC-content. This suggests that the annealing of free floating adapters present in the sequencing libraries does not occur randomly. The underlying mechanisms are not yet well understood but could be related to differences in the DNA denaturation temperatures between DNA fragments of different size. Due to the lower denaturation temperature, short fragments might be occurring at a higher rate in single-stranded conformation and are thereby more accessible to free floating index primers. Since shorter fragments in ancient DNA libraries often represent endogenous DNA, whereas longer fragments are mostly environmental contamination (Green *et al*. 2010), index-hopping can disproportionally affect the reads of interest in ancient DNA libraries.

To further illustrate the effect of index hopping on estimates of endogenous content and population genomics inferences, we employed simulations that encompass the complete range of reported index hopping rates and span a distribution of endogenous content typical for ancient DNA studies. Through simulations, we show that biases due to misassigned reads start to appear at index hopping rates below 0.5% when analysing samples of low coverage (<3X). As samples with low endogenous content predominantly act as receivers of hopped reads, inferences of population differentiation become less clear (Figure 3c) due to many hopped reads being erroneously assigned to low quantity samples. This also results in the false inference of shared ancestry between individuals from divergent populations as exemplified by Admixture (Figure 3d). In contrast, the inference of gene flow between populations through D-statistics is relatively robust to the biases resulting from index-hopping, if the proportion of misassigned reads between the tested samples is similar (Figure S6). In these cases, both samples contain similar proportions of false alleles from the 3rd ingroup population and therefore no significant deviation from zero is observed. Nonetheless, if coverage between the two tested samples is highly different (and thus one of the samples has a higher proportion of misassigned reads), Z-scores become skewed and might be falsely interpreted as a signal of gene flow (Figure S6). At the rate of index-hopping reported here (~0.470%), only genomes above 3X coverage remain largely unbiased, and thus for ancient DNA studies where samples are being multiplexed, elimination of index-hopping is of great importance.

We show that even with a low rate of index-hopping, such as the one observed in our empirical study, downstream inferences can become biased if sample qualities are highly different. Therefore, variation in sample endogenous DNA content are ideally kept to a minimum when sequencing a sample pool on the same lane. Pre-pooling qPCR quantification of sample DNA (and endogenous content) can be helpful to balance the sequencing libraries. Additionally, when multiplexed samples are sequenced to high depth (i.e. across multiple lanes/flowcells), re-pooling could be considered after the first sequencing run if high variation in (endogenous) read numbers is observed. This is especially relevant for the NovaSeq platform, the most powerful sequencing platform currently available, since it has been specifically designed for the multiplexing of up to hundreds of samples.

We show that in cases where low coverage data is generated or absolutely certainty is required, even a low remaining rate of misassigned reads can cause severe downstream biases. For such studies we therefore recommend the use of either short barcoded in-line adapters and/or dual indexing when preparing pooled libraries for next generation sequencing, independently of sequencing platform.

## Author contributions

TvdV and MO performed the wetlab experiments. TvdV and FV performed computational data-analysis of the sequenced libraries. TvdV ran the simulations. TvdV and KG established the experimental design. TvdV, LD and KG conceived the study, interpreted the results and wrote the manuscript (with contribution from all authors).

### Acknowledgments

We acknowledge the support from the Science for Life Laboratory, the Knut and Alice Wallenberg Foundation, the National Genomics Infrastructure funded by the Swedish Research Council, and Uppsala Multidisciplinary Center for Advanced Computational Science for assistance with massively parallel sequencing and access to the UPPMAX computational infrastructure. We thank Illumina for providing sequencing reagents. Illumina had no role in study design, data collection and analysis, decision to publish or preparation of the manuscript.

## Data access

Raw sequencing data is available at the European nucleotide archive under accession number XXXX.

All script that were used to simulate the data are available on github: https://github.com/XXXX

## References

Besenbacher S, Hvilsom C, Marques-Bonet T, Mailund T, Schierup MH (2018) Direct estimation of mutations in great apes reveals significant recent human slowdown in the yearly mutation rate. bioRxiv, 287821.

Briggs AW, Stenzel U, Meyer M et al. (2010) Removal of deaminated cytosines and detection of in vivo methylation in ancient DNA. Nucleic Acids research, 38, e87.

Carlsen T, Aas AB, Lindner D et al. (2012) Don’t make a mista(g)ke: Is tag switching an overlooked source of error in amplicon pyrosequencing studies? Fungal Ecology, 5, 747–749.

Craig DW, Pearson J V, Szelinger S et al. (2008) Identification of genetic variants using bar-coded multiplexed sequencing. Nature Methods, 5, 887–893.

D’Amore R, Ijaz UZ, Schirmer M et al. (2016) A comprehensive benchmarking study of protocols and sequencing platforms for 16S rRNA community profiling. BMC genomics, 17, 55.

Esling P, Lejzerowicz F, Pawlowski J (2015) Accurate multiplexing and filtering for high-throughput amplicon-sequencing. Nucleic Acids Research, 43, 2513–2524.

Fumagalli M, Vieira FG, Korneliussen TS et al. (2013) Quantifying population genetic differentiation from next-generation sequencing data. Genetics, 195, 979–992.

Gordon D, Huddleston J, Chaisson MJP et al. (2016) Long-read sequence assembly of the gorilla genome. Science, 352, aae0344–aae0344.

Green RE, Krause J, Briggs AW et al. (2010) A Draft Sequence of the Neandertal Genome. Science, 328, 710–722.

Griffiths JA, Richard AC, Bach K, Lun ATL, Marioni JC (2018) Detection and removal of barcode swapping in single-cell RNA-seq data. Nature Communications, 9, 2667.

Illumina Inc. NovaSeq 6000 Sequencing system. Illumina Inc. (2017) Effects of Index Misassignment on Multiplexing and Downstream Analysis.

Kircher M, Sawyer S, Meyer M (2012) Double indexing overcomes inaccuracies in multiplex sequencing on the Illumina platform. Nucleic Acids research, 40, e3.

Korneliussen TS, Albrechtsen A, Nielsen R (2014) ANGSD: Analysis of Next Generation Sequencing Data. BMC Bioinformatics, 15, 356.

Li H, Durbin R (2009) Fast and accurate short read alignment with Burrows-Wheeler transform. Bioinformatics, 25, 1754–1760.

Meyer M, Kircher M (2010) Illumina sequencing library preparation for highly multiplexed target capture and sequencing. Cold Spring Harbor Protocols, 5, pdb.prot5448.

Meyerhans A, Vartanian JP, Wain-Hobson S (1990) DNA recombination during PCR. Nucleic Acids Research, 18, 1687–1691.

Mitra A, Skrzypczak M, Ginalski K, Rowicka M (2015) Strategies for achieving high sequencing accuracy for low diversity samples and avoiding sample bleeding using Illumina platform (C Oudejans, Ed,). PLOS ONE, 10, e0120520.

Nelson MC, Morrison HG, Benjamino J, Grim SL, Graf J (2014) Analysis, Optimization and Verification of Illumina-Generated 16S rRNA Gene Amplicon Surveys (MM Heimesaat, Ed,). PLOS ONE, 9, e94249.

Odelberg SJ, Weiss RB, Hata A, White R (1995) Template-switching during DNA synthesis by Thermus aquaticus DNA polymerase I. Nucleic Acids research, 23, 2049–57.

Owens GL, Todesco M, Drummond EBM, Yeaman S, Rieseberg LH (2018) A novel post hoc method for detecting index switching finds no evidence for increased switching on the Illumina HiSeq X. Molecular Ecology Resources, 18, 169–175.

Renaud G, Stenzel U, Maricic T, Wiebe V, Kelso J (2015) deML: robust demultiplexing of Illumina sequences using a likelihood-based approach. Bioinformatics (Oxford, England), 31, 770–2.

Rohland N, Harney E, Mallick S, Nordenfelt S, Reich D (2015) Partial uracil-DNA-glycosylase treatment for screening of ancient DNA. Philosophical transactions of the Royal Society of London. Series B, Biological sciences, 370, 20130624.

Rohland N, Reich D (2012) Cost-effective, high-throughput DNA sequencing libraries for multiplexed target capture. Genome Research, 22, 939–946.

Schmieder R, Edwards R (2011) Quality control and preprocessing of metagenomic datasets. Bioinformatics, 27, 863–864.

Schubert M, Lindgreen S, Orlando L (2016) AdapterRemoval v2: rapid adapter trimming, identification, and read merging. BMC Research Notes, 9, 88.

Shapiro B, Hofreiter M (2014) A paleogenomic perspective on evolution and gene function: New insights from ancient DNA. Science, 343, 1236573–1236573.

Sinha R, Stanley G, Gulati GS et al. (2017) Index Switching Causes “Spreading-Of-Signal” Among Multiplexed Samples In Illumina HiSeq 4000 DNA Sequencing. bioRxiv.

Skoglund P, Northoff BH, Shunkov M V et al. (2014) Separating endogenous ancient DNA from modern day contamination in a Siberian Neandertal. Proceedings of the National Academy of Sciences of the United States of America, 111, 2229–34.

Skotte L, Korneliussen TS, Albrechtsen A (2013) Estimating Individual Admixture Proportions from Next Generation Sequencing Data. Genetics, 195, 693–702.

Smith AM, Heisler LE, St.Onge RP et al. (2010) Highly-multiplexed barcode sequencing: an efficient method for parallel analysis of pooled samples. Nucleic Acids Research, 38, e142–e142.

Team R Core (2016) R: A language and environment for statistical computing. R Foundation for Statistical Computing, Vienna, Austria. ISBN 3-900051-07-0, URL http://www.R-project.org/.

van der Valk T, Sandoval-Castellanos E, Caillaud D et al. (2018) Significant loss of mitochondrial diversity within the last century due to extinction of peripheral populations in eastern gorillas. Scientific Reports, 8, 6551.

Vodák D, Lorenz S, Nakken S et al. (2018) Sample-Index Misassignment Impacts Tumour Exome Sequencing. Scientific Reports, 8, 5307.

Wright ES, Vetsigian KH (2016) Quality filtering of Illumina index reads mitigates sample cross-talk. BMC genomics, 17, 876.

